# The Protein Hourglass: Exact First Passage Time Distributions for Protein Thresholds

**DOI:** 10.1101/2020.08.14.251223

**Authors:** Krishna Rijal, Ashok Prasad, Dibyendu Das

**Affiliations:** Physics Department, Indian Institute of Technology Bombay, Mumbai 400076, India; Department of Chemical and Biological Engineering, Colorado State University, Fort Collins, Colorado, USA

## Abstract

Protein thresholds have been shown to act as an ancient timekeeping device, such as in the time to lysis of *E. coli* infected with bacteriophage *lambda*. The time taken for protein levels to reach a particular threshold for the first time is defined as the first passage time of the protein synthesis system, which is a stochastic quantity. The first few moments of the distribution of first passage times were known earlier, but an analytical expression for the full distribution was not available. In this work, we derive an analytical expression for the first passage times for a long-lived protein. This expression allows us to calculate the full distribution not only for cases of no self-regulation, but also for both positive and negative self-regulation of the threshold protein. We show that the shape of the distribution matches previous experimental data on lambda-phage lysis time distributions. We also provide analytical expressions for the FPT distribution with non-zero degradation in Laplace space. Furthermore, we study the noise in the precision of the first passage times described by coefficient of variation (CV) of the distribution as a function of the protein threshold value. We show that under conditions of positive self-regulation, the CV declines monotonically with increasing protein threshold, while under conditions of linear negative self-regulation, there is an optimal protein threshold that minimizes the noise in the first passage times.

## I. INTRODUCTION

Thresholds in protein concentrations are ubiquitous in cell biology and are thought to be one of the main methods by which cells control the timings of events. Imposing a threshold before a biological process can be activated not only protects the process from noise-driven spurious activation, it also allows the cell to maintain a certain time interval between events. Cells do have other timing devices, most significant of which are circadian and ultradian rhythms. But thresholds offer one of the simplest timing devices possible, like the sand-filled hourglass of yesteryears, which is probably why they are ubiquitous in cellular processes. A good example of thresholds determining timing is provided by the passage of cells through the restriction point (R) during the cell cycle. It has been shown that passage through R in fibroblasts is governed by a rather precise threshold of the activity of the kinase, Cdk, and that a live cell Cdk sensor accurately predicts passage through R for 96% of cells [1]. Just as in cell reproduction, in cell death too, the timing and probability of mitochondrial Outer Membrane Permeabilization, the irreversible step committing the cell to apoptosis after an event like DNA damage, depends on pro-apoptotic signals crossing a threshold[2–4]. However, the most well studied question in cell biology where thresholds are believed to play a major role in determining timing is the replication of lambda-phage.

After lambda-phage infects an E. coli bacterium, it can have one of two possible life cycles, either the lytic or the lysogenic path. The molecular basis of this decision making has been critical to the development of our understanding of stochastic gene regulation over the last half-century[5–15]. The lysogenic phage incorporates their DNA into the bacterial chromosome, which sits there silently until a stress signal indicating host distress activates the genes and they enter the lytic cycle. In the lytic pathway, the virus multiplies in its host for a programmed time period, and then lyses the host, which then bursts, releasing a number of virions. The majority of infected bacteria will take the lytic route, and most remarkably, the timing of lysis after initial phage infection is tightly controlled [16] with a variation of only about 5% of the mean, which is a remarkable level of precision for such a complicated system, especially when the burst size is found to vary significantly more [16–18]. The timing system of lambda-phage is controlled by the accumulation of a protein, holin [19, 20], that permeabilizes the membrane after it accumulates to a critical level. The tight regulation of lysis time is believed to have an evolutionary basis and was a prediction from optimal foraging theory[16]. Significantly there does not appear to be any additional genetic regulation that determines the timing of lysis. In two seminal papers [21, 22], Singh, Dennehy and colleagues explored the idea that the timing of lysis is controlled by the first passage time for the holin protein to reach a threshold. For any stochastic system, the first passage time is a random variable that denotes the first time some event occurred [23]. Examples of such events may be like a passive or active particle reaching a boundary [24, 25], a protein binding to a specific patch on DNA [26], a moving kinetochore being captured by microtubules [27], or as in this case protein levels reaching a pre-determined threshold. The papers [21, 22], showed that a model based on the statistics of first passage times to reach a critical protein concentration could reproduce the observed data. Furthermore they provided a basis for the observed lack of regulation of the timing by demonstrating that for a long-lived protein, the optimal strategy for precise event timing is no regulation at all, and that both negative and positive feedback actually increase the noise in event timing [22]. The above-mentioned papers were based on calculations of the first few moments of the first passage time distribution, while an analytical expression of the full distribution itself was not available prior to this paper.

There are significant advantages in obtaining the full distribution of a stochastic biochemical process. For example, it has been shown that when inferring parameters from data, it is possible to infer incorrect parameters with only the mean and the variance of the distribution [28]. However, given the complexity in even simple molecular biological processes, there are relatively few exact distributions known. These include Refs. [29–35]. In this paper we present an analytical calculation for the first passage time of a long-lived protein driven by a bursting process to reach a threshold. We show that the first passage distribution applies even when the protein has negative or positive autoregulation, showing either linear kinetics or Hill kinetics. We show that the shape of the distribution matches the experimental data on lysis times. We calculate the coefficient of variation of the distribution and show how it varies with the level of the protein threshold. Finally, for non-zero protein degradation, we calculate the Laplace transform of the FPT distribution exactly.

## II. MODEL

Consider a gene that is switched on at time *t* = 0 and begins to express a timekeeper protein, which triggers an intracellular event of interest, eg. lysis, once it reaches a critical level in the cell. A minimal model of gene expression assumes translation in bursts and can incorporate feedback regulation by considering the transcription rate as a function of the protein level (see Fig.(1)). We represent the protein population count or number by states [0, 1, *…, X*]. Protein numbers go up due to bursts of transcription-translation that add a number of the proteins to the system, while protein numbers are reduced due to degradation of single proteins. The process terminates when the protein level reaches a certain threshold or absorbing-state *X* for the first time. The corresponding time *t* defines the “first passage time” (FPT). The statistics of these times determines the timing of the intracellular event.

**FIG. 1:**
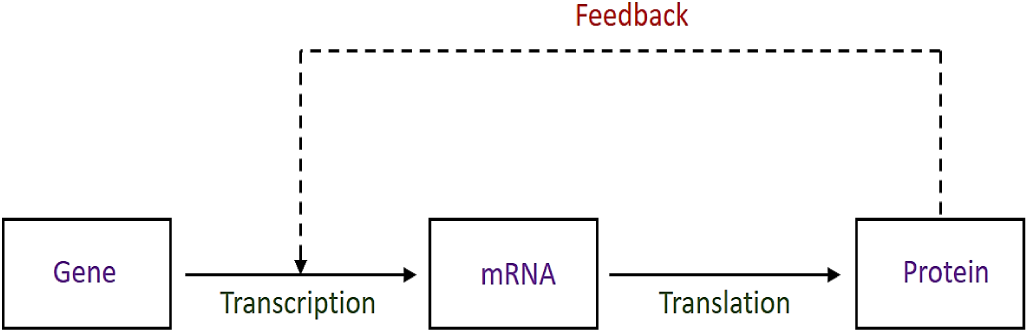
Model depicting a gene transcribing mRNAs, which are further translated into proteins. The transcription rate is assumed to be regulated by the protein level, creating a feedback loop (denoted by the dashed line).

The bursts of protein numbers are assumed to follow a geometric distribution, as shown previously for protein burst-size distributions [36]. Thus the Poisson rate of making a transition from state *i* to state *i* + *n* is

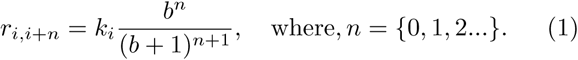

Here mean burst size is denoted by *b ∈* (0, *∞*), and *n* = 0 denotes no change in protein number with probability *b/*(*b*+1). The net Poisson rate for any burst of size *n >* 0 out of state *i* is thus

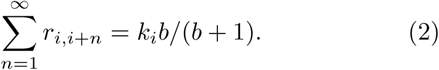

Each protein molecule degrades with a constant rate *γ*. For most of our calculations, we will assume that the proteins are long-lived and hence we will set the degradation rate *γ* to 0.

We use the backward Master equation (BME) formalism [37] to calculate the first passage probabilities. The BME describes the time-evolution of the survival probabilities *S*(*i, t*), that we define as the probability that the protein count stays below the threshold *X* at time *t*, given that it was at *i* at time *t* = 0, where *i* = 0, 1, *…*, (*X −*1)).

If we define a vector **S**(*t*)= [*S*(0, *t*) *S*(1, *t*) *S*(2, *t*) *… S*(*X −* 1, *t*)]^*T*^ whose components are survival probabilities, we may write the backward master equation as

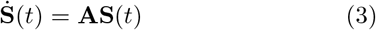

with the initial condition **S**(*t* = 0) = [1 1 1 *…* 1]^*T*^. Since the process terminates for protein count reaching any value *≥ X*, the boundary condition is *S*(*i ≥ X, t*) = 0. Taking into account degradation, the above matrix of Poisson rates, **A** is

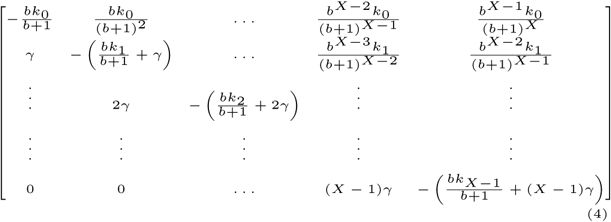

Note that the general element in this rate matrix may be written as

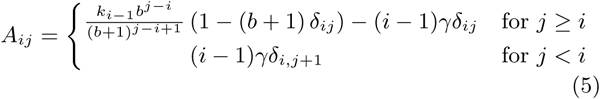

Furthermore, the diagonal elements of this matrix are the negative sum of all outgoing rates 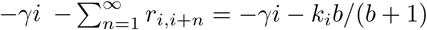.

The first passage probability to reach *X* for the first time between *t* to *t* + *dt* is *f*_0,*X*_ (*t*)*dt*, where the FPT distribution *f*_0,*X*_ (*t*) is related to the survival probability as *f*_0,*X*_ (*t*) = *− ∂S*_0,*X*_ (*t*)*/∂t*. Thus our objective is to solve for *S*_0,*X*_ (*t*) and obtain the desired expression for *f*_0,*X*_ (*t*).

From here, we can proceed using two different methods. Integrating Eq. (3) gives

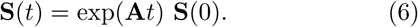

The FPT distribution follows from Eqs (3) and (6) as

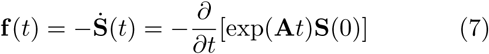

Calculation of the full distribution requires the evaluation of exp(**A***t*) which may be carried out as shown in Appendix G. An alternative and more elegant method is to use Laplace Transforms on Eq. (3) as outlined below. Both the methods to calculate the full distribution gives identical results as expected. Furthermore, we can also calculate the moments of the FPT through evaluation of the inverse matrix **A**^*−***1**^(as was also shown in [22]).

## III. RESULTS

### A. The exact first passage time distribution

#### 1. Unequal Transition Rates with zero protein degradation (γ = 0)

In the most general case the transition rate constants *k*_*i*_ are all unequal. We outline the solution in the case of *γ* = 0 as follows. We can take Laplace transform of Eq. (3), to convert it into an algebraic equation. If we define 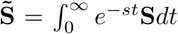, then Eq (3), after performing Laplace transformation gives,

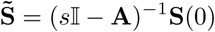

The general form of the elements of (*s𝕀 −* **A**)^*−*1^ is

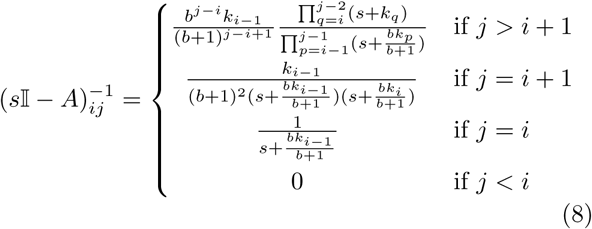

In order to obtain this result, we first evaluated (*s𝕀 −* **A**)^*−*1^ in the special cases of *X* = 1, 2, 3 and 4, and then generalized the mathematical form to any *X*. We are interested in the case of initial protein number being 0, and in the corresponding first passage distribution

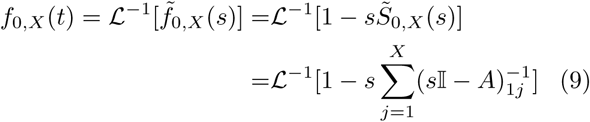

where,ℒ^*−*1^ denotes inverse Laplace transformation. In the last step, we have used the fact that **S**(*t* = 0) = [1 1 1 *…* 1] ^*T*^. Since all the poles of (*s𝕀 −* **A**)^*−*1^are simple, it is easy to find inverse Laplace transform by evaluating Bromwich integral, by finding residues. We get the exact FPT distribution (see Appendix A) as

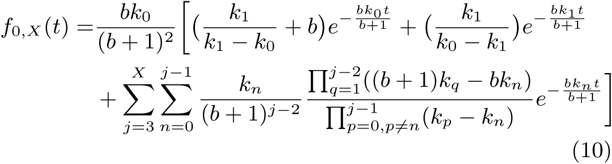

#### 2. Equal Transition Rates with zero protein degradation (γ = 0)

The case of uniform rates *k*_*i*_ = *k* (corresponding to no feedback) is special. We see in Eqs. (10) that denominators of various terms go to zero. This may appear to lead to divergent answers, which is not expected physically for any bounded function *k*_*i*_. From Eq. (8) setting all *k*_*i*_ = *k*, we get

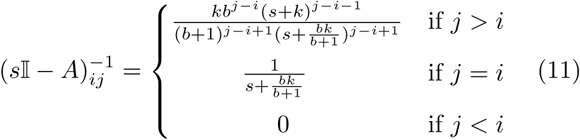

One way to verify Eq. (11) directly is to check that (*s𝕀 −* **A**)^*−*1^(*s𝕀 −* **A**) = *𝕀* (see Appendix B). Compared to Eq. (8), in Eq. (11) the poles are not simple but of higher order. Hence, the Laplace inverse transform leading to the FPT distribution will not be a sum of pure exponentials. We use Bromwich integral and find residues (see Appendix C). The final desired answer for the first passage time distribution is:

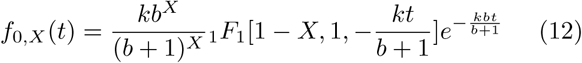

Here _1_*F*_1_ is the confluent hypergeometric function. An alternative derivation of Eq. (12), involving interesting mathematical identities, are shown in Appendix H and I.

#### 3. Non-zero protein degradation (γ ≠ 0)

The main results derived above are for zero protein degradation rates. However we have also obtained exact results in the Laplace space for non-zero degradation rates. Using Eq. (9), the result for general *X* may be obtained by studying the pattern of 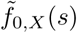 for small *X* values and then generalizing to higher values.

##### Unequal transition rates

For distinct *k*_*i*_ for *X >* 2, we have

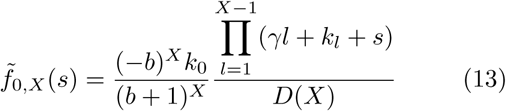

where the denominator *D*(*X*) of Eq. (13) is not easy to find in a compact form. But we could relate *D*(*X*) to *D*(*X* − 1) and another function *Q*(*X*), through a recurrence relation given below:

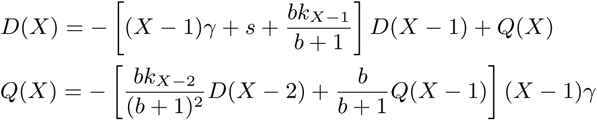

with 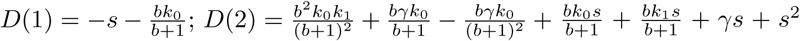. 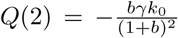and Using the recurrence relations one can iteratively obtain *D*(*X*) for any *X* and hence find the Laplace transform 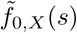.

##### Equal transition rates

For *k*_*i*_ = *k* for all *i*, the FPT distribution in the Laplace space is simpler. For

X > 1,

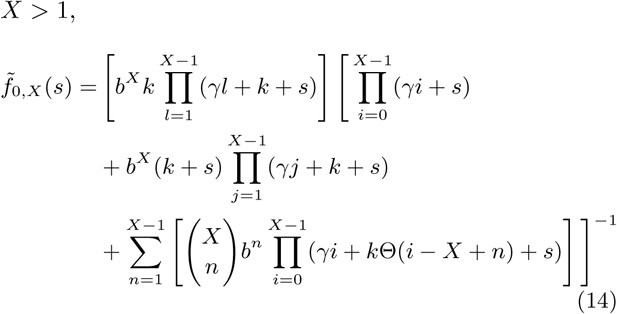

where Θ denotes Heaviside step function.

In both the cases of unequal and equal transition rates, the expressions of 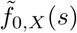 have polynomials of degree equal to *X* as denominators (see Eq. (13) and (14)). It is well known that finding roots exactly of a polynomial beyond some small values of *X* is very difficult. Hence, finding inverse Laplace transform for large *X* remains intractable for *γ ≠* 0. To demonstrate the complexity of the poles for even small values of *X*, we have calculated the poles and hence *f*_0,*X*_ (*t*) for *X* = 1, 2, 3 in Appendix D and E, for both the cases.

### B. Kinetic Monte Carlo results compared with the exact formulae

In order to verify Eq. (10) and Eq. (12) we carried out Kinetic Monte Carlo (KMC) simulations for cases of positive, negative and no feedback. In order to compare relative fluctuations, we hold the mean first passage time same in the three cases. The FPT distribution is obtained by measuring the times of first passage to reach the protein number threshold of *X* = 500 starting from 0 number of proteins. At any instant of time, the system is in a state *i* with *i* number of proteins. In the next step of KMC, any one of the events by which *i → i* + *n* is permitted, where in principle *n ∈* [1, *∞*). But for practicality, to have a finite set of events in simulations, we restrict *n* ∈ [1, *X*]. The rates of the transition of *i → i*+*n* are given by Eq. (1). As is usual for KMC, the events are chosen with probability *r*_*i,i*+*n*_*/Q*_*i*_, and the time increments Δ*t* are drawn randomly from an exponential distribution with decay constant *Q*_*i*_, where *Q*_*i*_ is given by Eq. (2). For the simulations, following [22] we chose *X* = 500 protein molecules, and *k*_*i*_ = *c*_1_ *± c*_2_*i* for positive and negative feedback cases respectively. We have *c*_2_ = 0.05 min^*−*1^ for these two cases. For no feedback *c*_2_ = 0. The value of *c*_1_ was chosen to be 0.581 min^*−*1^, 25.443 min^*−*1^ and 6.275 min^*−*1^ for the positive, negative and zero feedback cases respectively, to ensure that all the three cases have a same mean first passage time of 40 mins (for ease of comparison). We have also chosen *b* = 2 just as in [22].

The plots of our KMC simulations and the curves for *f*_0,*X*_ (*t*) given by the exact theoretical formulae Eqs. (10) and (12) are shown in Fig. (2), and they match perfectly.

**FIG. 2:**
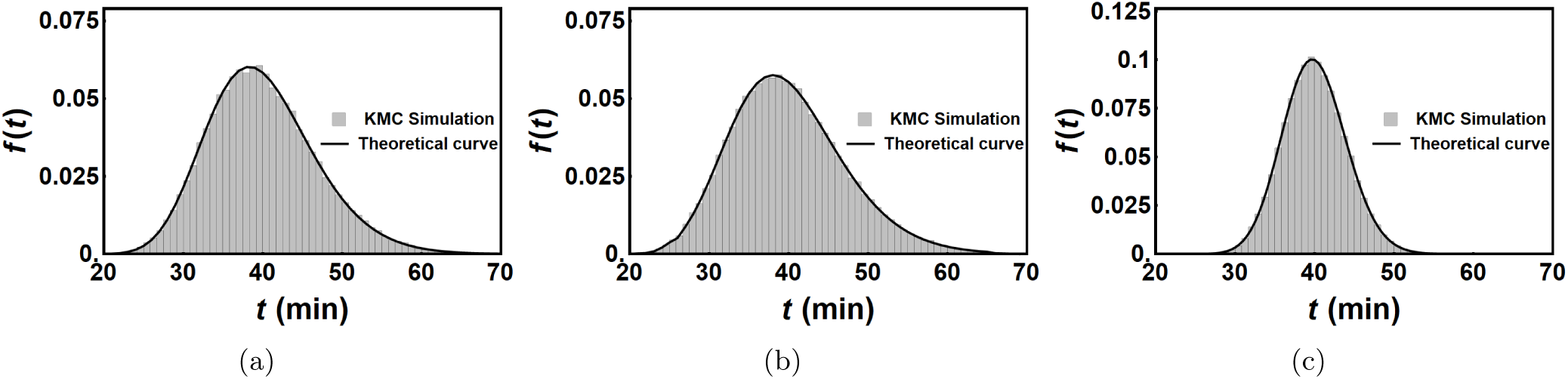
Theoretical FPT distributions *f* (*t*), are compared with KMC simulations. For stable protein (*γ* = 0), burst size *b* = 2, and protein threshold level *X* = 500. We show histogram plots of 100000 simulations performed for linear feedbacks of the for *k*_*i*_ = *c*_1_ + *c*_2_*i* with (a) *c*_2_ = *−*0.05 min^*−*1^ for positive feedback, (b) *c*_2_ = − 0.05 min^*−*1^ for negative feedback and (c) *c*_2_ = 0 for the no feedback cases. Value of *c*_1_ is choosen such that mean FPT distribution is kept constant at 40 min. The black curves on the top of the histograms are the theoretical FPT distributions given by Eq. (10) and Eq. (12), for the chosen parameters.

### C. Formulae for moments and cumulants

We can also calculate exact expressions for the moments of the FPT distribution, using Eq. (7) after obtaining the matrix **A**^*−*1^. The general form of elements of the matrix **A**^*−*1^ is

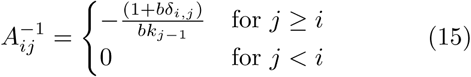

We obtained this by putting *s* = 0 in Eq. (8). Identical expression was also obtained in [22]. For special case of all identical *k*_*i*_ = *k*, first two moments are given as [22]

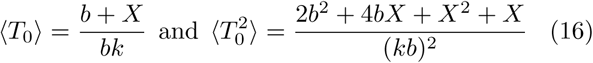

which gives

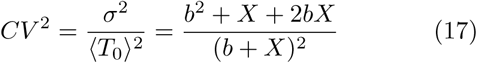

But the explicit forms of the higher moments and cumulants were not given in [22]. Here, for the ready reference of the reader we are giving the explicit expressions for the next two higher cumulants (see calculations in Appendix F):

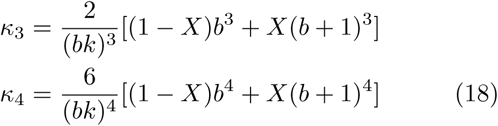

The following definitions of skewness and kurtosis based on *κ*_3_ and *κ*_4_ will be used in the next section:

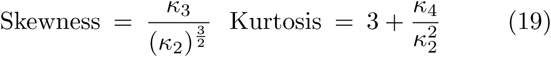

### D. Experimental Lysis Time Distribution in Lambda phage compared to the Theoretical Distribution

Lysis of the host E. coli by an infective lambda-phage is controlled by a protein from the holin family, S105 or simply holin, that accumulates in the bacterial cytoplasmic membrane till it reaches a critical value, when it triggers to form micron-scale holes in the membrane, releasing an enzyme that rapidly destroys the bacterial cell wall and frees the viral progeny. The timing of lysis has been shown to be tightly and precisely controlled by the accumulation of the holin protein, i.e. by a first passage time process. Since holins have a half-life larger than the mean lysis time, our results for the FPT of a long-lived protein are directly applicable [17, 20–22, 38, 39]

Holin synthesis does not begin immediately after infection by the lambda phage, but after a delay. In order to compare the theoretical distribution with the experimental data, we first have to account for this delay in the transcription of holin, *T*_*Pr*_*′*, which is known from experiments [21] to be *≈* 15 mins. We will also assume, following [21], that the standard deviation of *T*_*pR*_*′* is small enough to ignore. So lysis time *t*_*LT*_ can be related to first passage time *t* as:

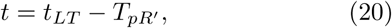

and it follows that

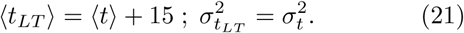

We now make the simplified assumptions, following [21, 22], that: (i) there is no feedback, i.e. *k*_*I*_ = *k*, and (ii) the holins do not degrade easily, i.e. *γ* = 0. Experimentally [21] it is known that ⟨*t*_*LT*_⟩ *≈* 65 min, and *σ*_*LT*_ *≈* 3.5 min. Just as in [21], we take *X* = 1500. These numbers may be used in Eqs. (16) and (17) to obtain the the parameters

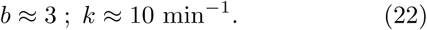

Finally using the above parameters (Eq. (22)), we plot the theoretical curve for *F* (*t*_*LT*_) (Eq. (12)) in Fig. (4) against the available experimental data from [21]. We can see that the theoretical distribution, with no fitting but using the parameters from [22] matches the experimental data quite well.

We tested to see whether we can improve the correspondence by fitting the parameters to the data. We fixed the values of mean first passage time to the experimentally determined 65 mins, and that of the burst size *b* to be 3. Using minimization of the residual sum of squares [40] we found that the best-fit value of X to be very close to 1500, namely 1463. Next, we kept only the mean fixed at 65 min and performed a 2-parameter minimization of the residual sum of squares. The best-fit value of *b* and *X* was found to be 2.6 and 1296 respectively. However the quality of the fits are not very significantly better than the original set of parameter values.

Using *b* = 3; *X* = 1500; *k* = 10 min^*−*1^ we can also compare the skewness and kurtosis of our predictions with the experimental values. The theoretical expressions are shown in Eq. (19) in the previous section. The correspondence is very close — the theoretically predicted skewness and kurtosis are 0.0025 and 3.015 as compared to the experimental values 0.0012 and 3.540, respectively.

### E. Feedback Effects

Eq. (10) is couched in terms of generic *k*_*i*_, which is the parameter that controls protein synthesis rates in the model. As can be seen from Eq. (2), *k*_*i*_ controls the Poisson rate out of state *i*. In general *k*_*i*_ can be a function of *i*, especially if the protein production is under some form of auto-regulation, as is very common in protein synthesis. Thus *k*_*i*_ could be constant, increase, or decrease as *i* increases, which corresponds to the cases of no regulation, positive regulation and negative regulation respectively. For both positive and negative self-regulation, we can numerically calculate the full distribution of first passage times by inputting the form of the self-regulation in Eq. (10). One of the important questions we can ask is whether incorporation of feedback has an impact on the noise in first passage times or not, which determines the accuracy of a FPT-based molecular clock. For a given level of threshold, [22] already compared the relative noise for the case of no feedback, positive feedback and negative feedback and found that the no feedback case typically has the lower level of noise. We show in Eq. (17) that for the no-regulation case the CV depends only on the burst size and the value of the threshold *X*. With addition of any kind of feedback, negative or positive, we are adding extra noise as a consequence of temporal inhomogeneity of the translation process, on top of the unavoidable Poissonian noise of translational burst.

We can also examine the behavior of the noise in the FPT distribution as a function of the protein threshold, *X*. We do this below for the cases of positive and negative feedback.

#### 1. Positive Feedback

We implemented positive feedback using either a linear function,

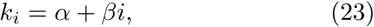

or a Hill function,

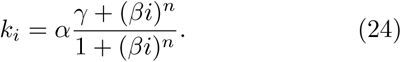

For numerical calculation we need to choose explicit numerical forms of these functions. Using arbitrarily chosen numerical parameters we plotted the coefficient of variation of the distribution as calculated using our results above against the protein threshold, *X*. The results are shown in Fig. 5, and they show that the relative noise in the FPT distribution reduces with increasing *X* (or equivalently with increasing mean FPT), but then effectively plateaus. Interestingly this behavior can be seen in both forms of the feedback.

**FIG. 3:**
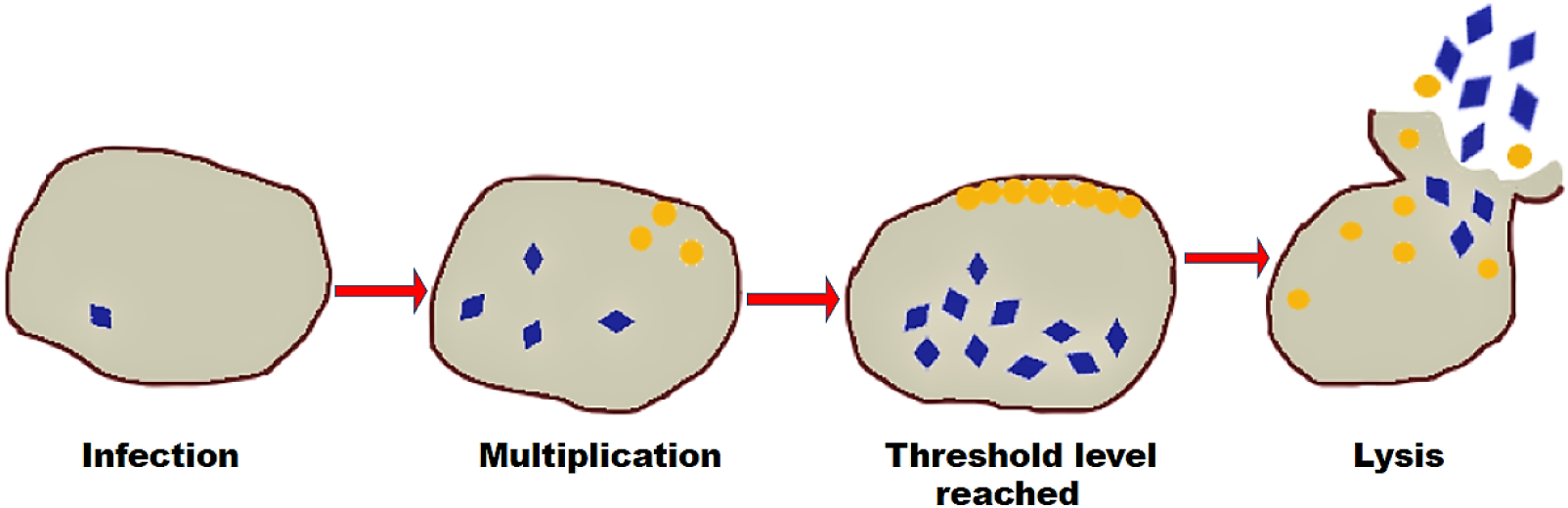
A schematic figure of steps leading to lysis of an E.Coli cell infected by a *λ*-phage. The blue rhombi represent viruses (in capsids), and the yellow circles represent the holin protein molecules.

**FIG. 4:**
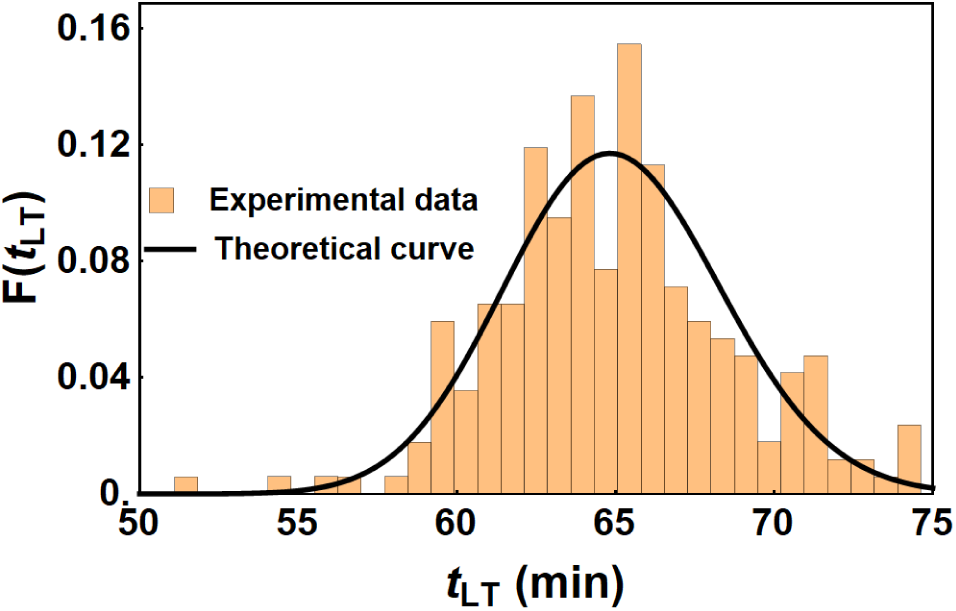
Comparison of experimental [21] and theoretical lysis time distributions. Here we use *b* = 3, *k* = 10 min^*−*1^ and *X* = 1500 for the theoretical formula (Eq. (12)).

**FIG. 5:**
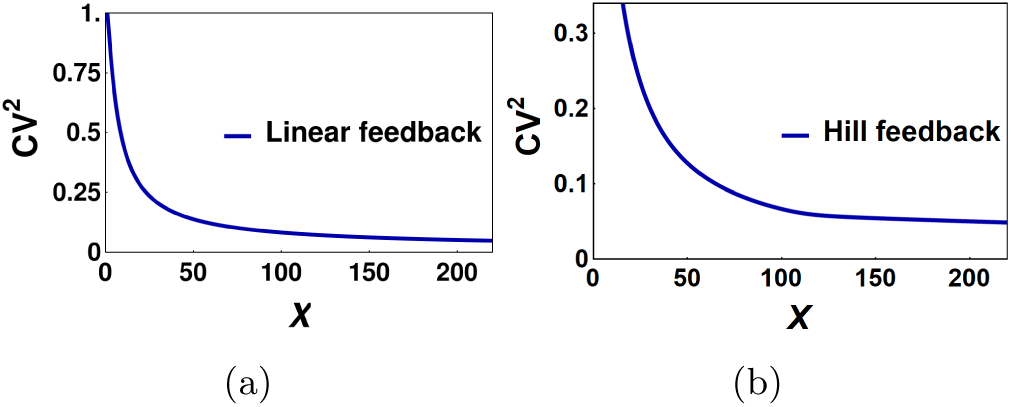
For stable protein (*γ* = 0) and for burst size, *b* = 3, noise in FPT (*CV* ^2^) is plotted as a function of protein threshold value (*X*) for different kinds of positive feedback: (a) Linear feedback, *k*_*i*_ = 1.5 + 0.0725*i* and (b) hill feedback 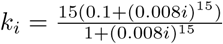

#### 2. Negative Feedback

The case of negative feedback turned out to be more interesting. We used a linear form of negative feedback, *k*_*i*_ = *α − βi*, as well as a Hill function form, given by

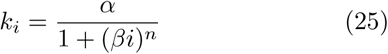

Our results are shown in Fig. 6. It may be seen that for the Hill function there is a monotonic decrease, while for the linear case, the noise in the FPT times first decreases and then increases, yielding a U-shaped curve. It was recently shown in [41] that the CV of lysis timing for different bacteriophage mutants as a function of the effective protein threshold is also U-shaped. In that paper they attributed this effect to the effect of dilution due to bacterial growth. We show here that a general negative feedback that may arise due to dilution, or possibly could be caused by some as yet undiscovered mechanism, reproduces this relationship.

**FIG. 6:**
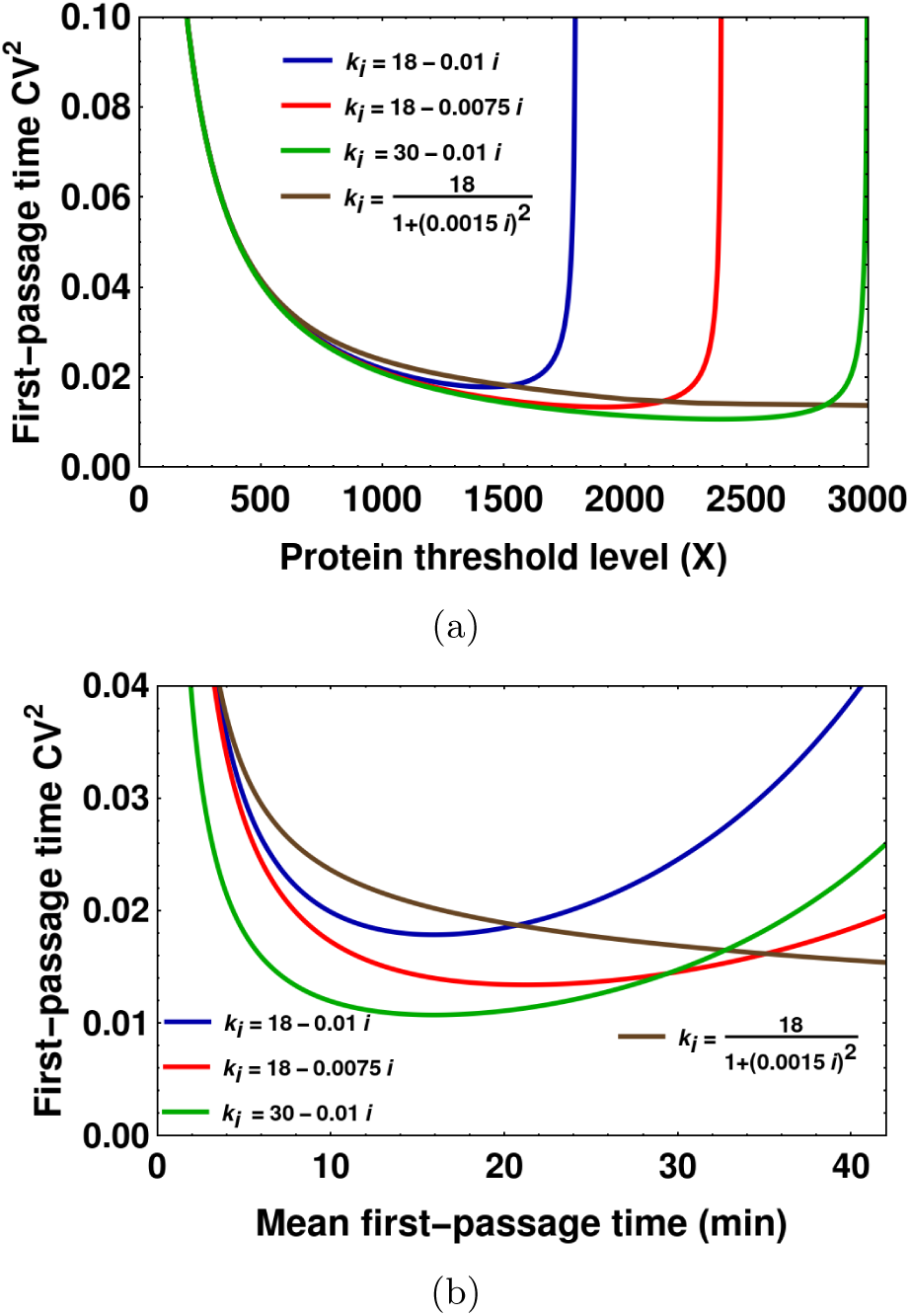
For stable protein (*γ* = 0) and for burst size, *b* = 10, noise in FPT (*CV* ^2^) is plotted as a function of (a) protein threshold value (*X*) and (b) mean FPT, for linear and hill negative feedbacks are plotted. For the linear feeback, the transription rate amplitude and feedback strengths are varied to see how the optimal mean and *X* depends on them.

## IV. DISCUSSION

In biology, it is often very difficult to find exact distributions of even simple stochastic processes. In those cases, people resort to expressions of mean and variance. But, the mean and variance contain partial information about a process, and in particular do not capture large deviations from central values. Analytic expression for full distributions are thus desirable when possible.

Many a time, the differences in time scales of various sub-processes involved are exploited to simplify the dynamics of the stochastic process, in order to make the problem analytically solvable or simpler. For long time, people relied on mean and variance of protein concentration in the gene expression of 2-state promoter system. Only recently, for some limiting timescale of protein lifetime, the steady state protein distribution could be calculated for the same problem [29]. In another example,[35], time-dependent distributions of mRNA and protein numbers are obtained, for detailed model of gene expression, by using biologically relevant timescale approximations.

In this paper, we used a similar strategy, where in the bulk of the paper we ignored protein degradation, and calculated exact FPT distribution for protein thresholds. This assumption is valid for stable proteins whose mean lifetime is much longer than the event timescale. Otherwise, our results are very general. An additional strength of these results lie in being able to predict the exact distribution with arbitrary types of feedback. Many proteins exhibit auto-regulation, thus acting as self-activators or self-repressors, which could in principle manifest as arbitrary functional forms of feedback. Indirect feedback of protein levels upon their own transcription and translation is also common, and could again be approximated as self-regulation. In all such cases where we can impute a functional form for the parameter *k*(*i*), the exact FPT distribution can be calculated by inputting that form into the general formulae obtained above. In principle, we can also solve the inverse problem and get insight about the dynamics of the process by measuring the experimental FPT distribution and using it to deduce the functional form of the feedback. Thus high precision experimental data on the FPT of lysis by lambda phage or any other similar process would be very valuable to help us discriminate between these different possibilities using the analytical expressions derived here.

While the bulk of the paper operates under the assumption of long-lived proteins, we also calculate the exact FPT distribution in Laplace case for non-zero degradation. We show that for the case with protein degradation, the main analytical difficulty is the complexity of the poles of the Laplace Transform, which involve solutions of very high order polynomial equations. We have also studied the noise in first passage times as described by CV and shown that for linear negative feedback, it has a minimum at an optimal protein threshold as well as at a value of the mean first passage time.

Given their simplicity, protein hourglasses, where the cells time a process using the first passage time of a protein level reaching a threshold, are likely to be widespread in cell biology. One can exploit the analytic expression of the distribution we obtained, to find the probability distribution of other cellular processes that depend on the FPT. One such example is the stochastic number of viral progeny on lysis, or the viral burst-size distribution, after a host-cell infection. We hope to extend our work towards these applications in future.

## ACKNOWLEDGMENTS

D.D. would like to acknowledge SERB India (grant no. MTR/2019/000341) for financial support. A.P. would like to acknowledge hospitality and support from the IIT Bombay Physics department, where he was a visiting scholar for a month when some of this research was carried out. K.R. thanks IIT Bombay for Institute Ph.D. fellowship.

## Appendix A FPT distribution calculation for unequal transition rates and *γ* = 0

**Bromwich Integral**

For a Laplace transform *f* (*s*), the inverse transform is given by:

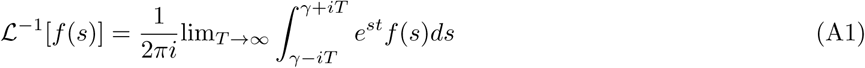

where the integration is done along the vertical line *Re*(*s*) = *γ* in the complex plane such that *γ* is greater than the real part of all singularities of *f* (*s*) and *f* (*s*) is bounded on the line. As we will see later, *f* (*s*) in our case is always of the form *P* (*s*)*/Q*(*s*), where *P* and *Q* are polynomials of *s* and degree of *Q* is greater than that of *P*. Such functions are always bounded over the entire complex plane except at the finite number of poles. So, we can always find the vertical line *Re*(*s*) = *γ* such that the boundedness condition is satisfied. Also,*P* (*s*)*/Q*(*s*) *→* 0 as *s → ∞*. So, we can do integration in an infinite D-shaped contour. Then, in our case, using residue theorem, Eq. (A1) becomes [42]

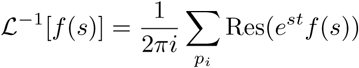

where, the summation is over all the poles, *p*_*i*_, of *f* (*s*). So,

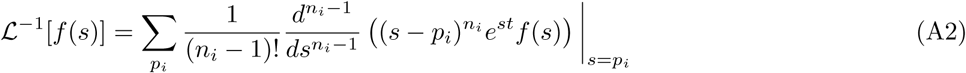

where, *n*_*i*_ is the order of *i*^*th*^ pole, *p*_*i*_.

For this general case of unequal transition rates with no protein degradation, recall from the main text

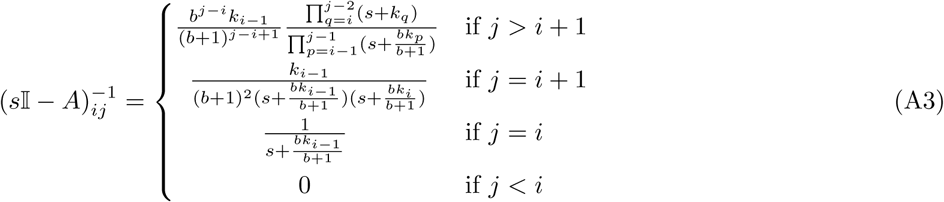

and

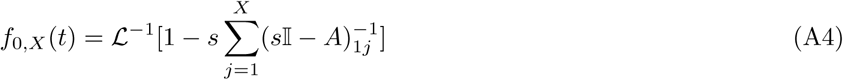

For *X* = 1

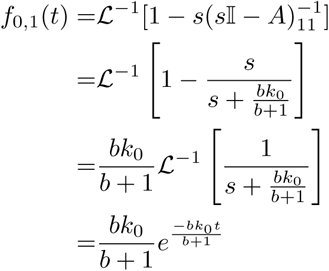

For *X* = 2

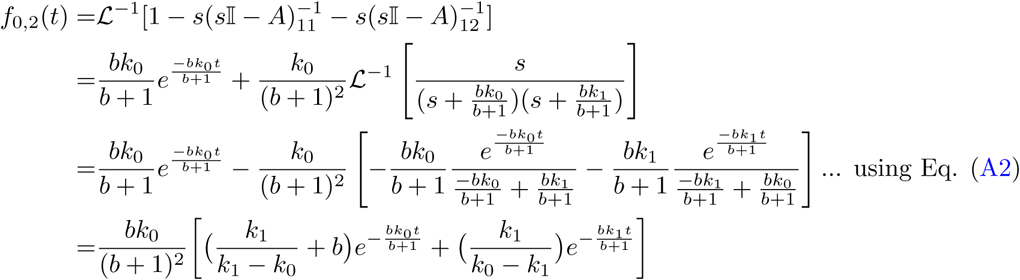

For *X* > 2

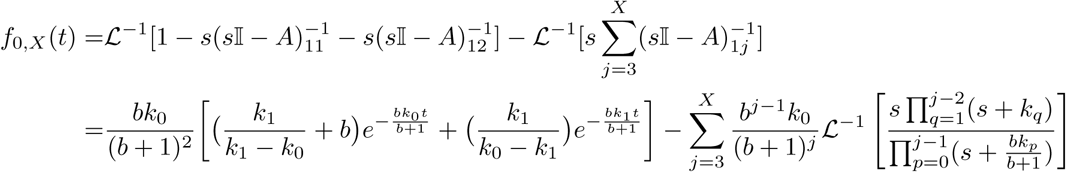

for each *j*, there are *j* number of simple poles at *bk*_*n*_*/*(*b* + 1) with *n* going from 0 to *j*1. So using Eq. (A2), we have

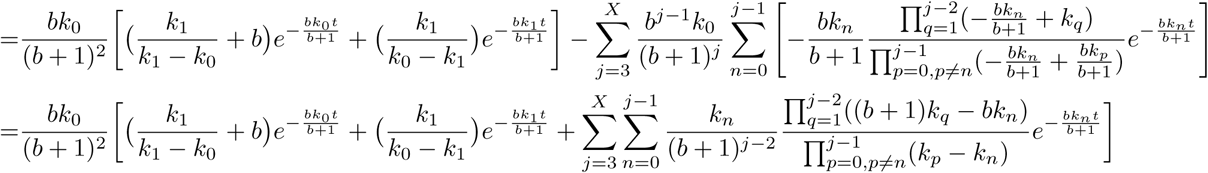

## Appendix B Proof of (*s*𝕀 *−* A)^*−*1^(*s*𝕀 *−* A) = 𝕀 for special case of equal transition rates and *γ* = 0

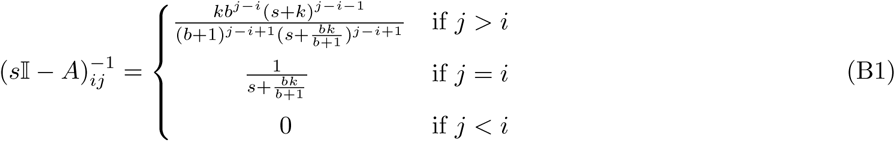

From Eq. (5) in main text,

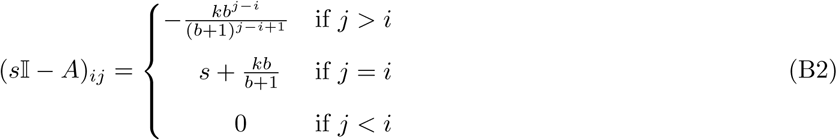

Product of two upper triangular matrices is also upper triangular. So, [(*s 𝕀 − A*)^*−*1^(*s 𝕀 − A*)]_*ij*_ = 0 for *j < i*.

Also, 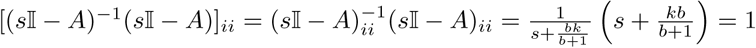

For *j* > *i*,

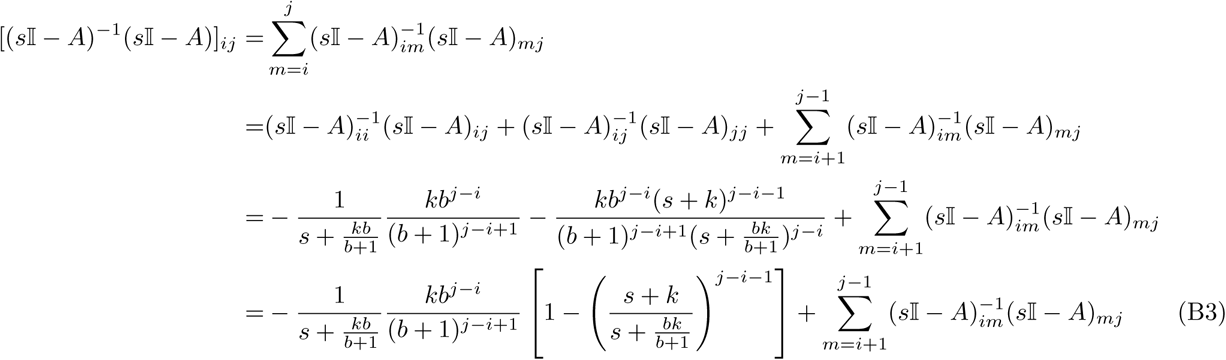

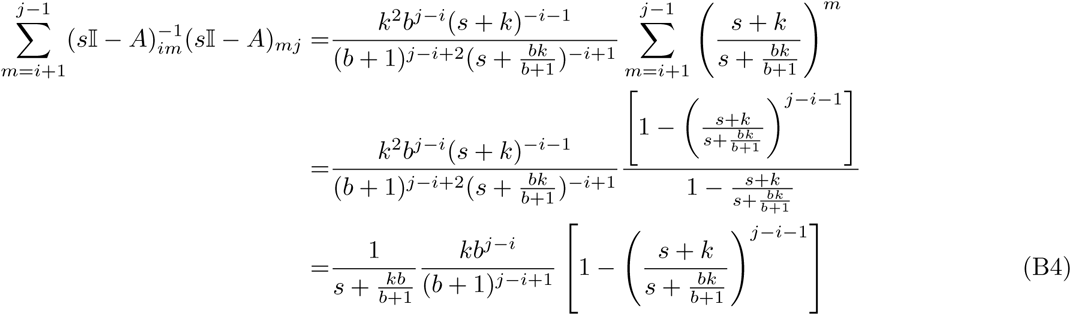

Putting Eq. (B4) into Eq. (B3), we get

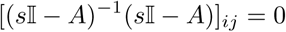

Hence, (*s 𝕀 −* **A**)^*−*1^(*s 𝕀 −* **A**) = *𝕀*

## Appendix C FPT distribution calculation for special case of equal transition rates and *γ* = 0

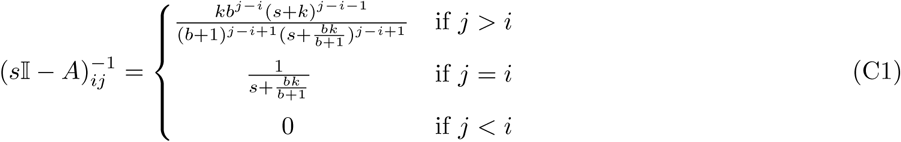

We know that 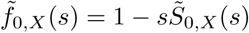. So,

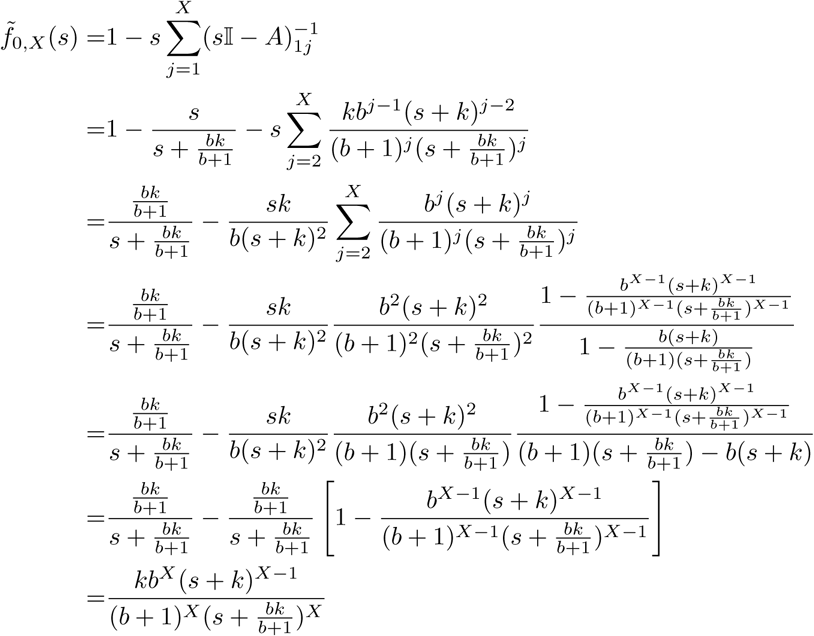

Using Eq. (A2),

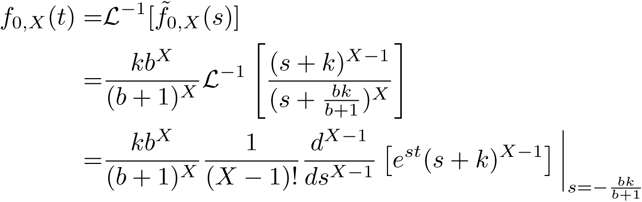

We can use Leibniz rule to calculate (*X −* 1)^*th*^ derivative

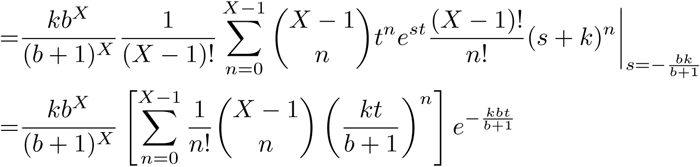

The term inside the square bracket is a confluent hypergeometric function _1_*F*_1_, and hence

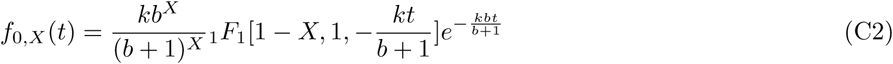

## Appendix D For small values of *X*, FPT distributions for unequal transition rates and *γ* ≠ 0

For *X* = 1, protein degradation is not relevant. So,

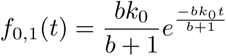

For *X* = 2,

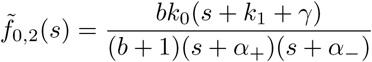

where

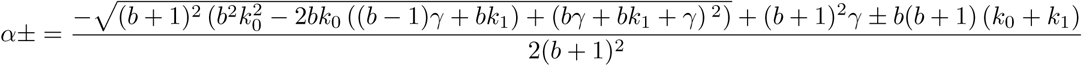

So, using Eq. (A2),

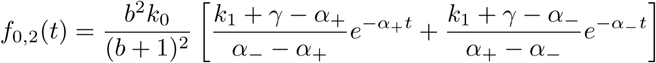

For *X* = 3,

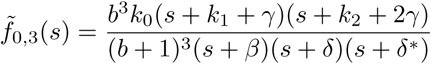

where *δ* and *δ*^*∗*^ are complex conjugates. The values of *β, δ* and *δ*^*∗*^ are too big to be put here. Using Eq. (A2), we get

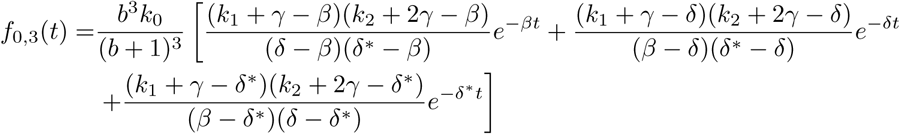

## Appendix E For small values of *X*, FPT distribution for equal transition rate and *γ* = 0

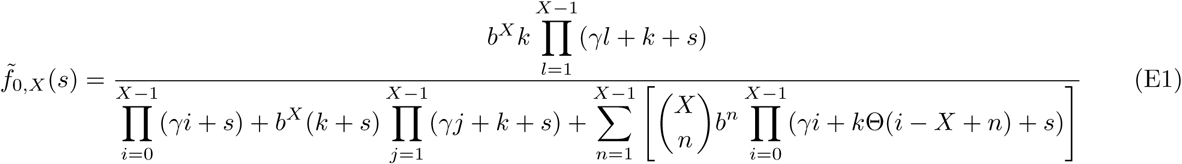

For *X* = 1, protein degradation is not relevant. So,

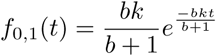

For *X* = 2,

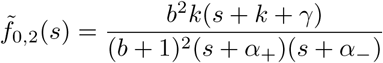

where 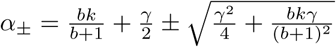

So, using Eq. (A2),

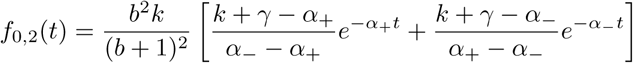

For *X* = 3,

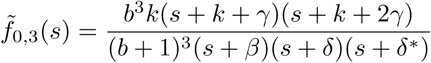

where *δ* and *δ*^*∗*^ are complex conjugates and

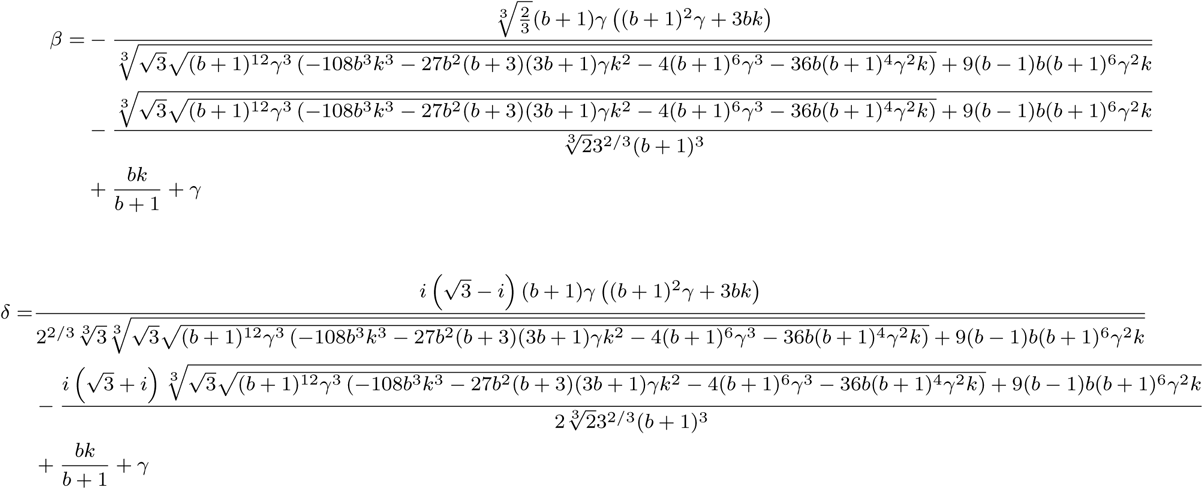

Using Eq. (A2), we get

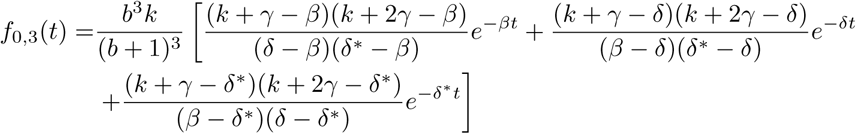

## Appendix F Moments calculation

We define a vector ⟨**T** ⟩^3^, whose components are third moments of the FPTs starting from various protein counts *i*. Doing integration by parts successively, we get:

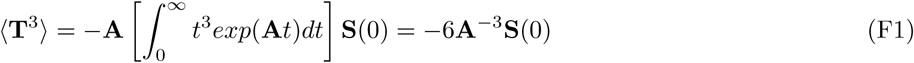

Noting the fact that

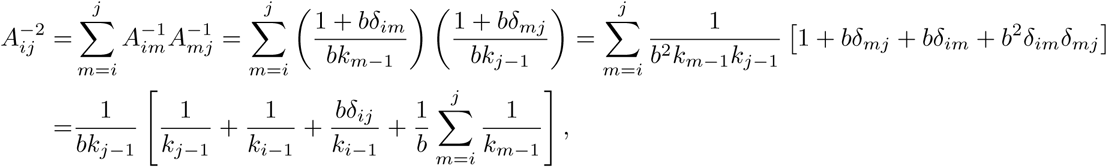

we finally have using Eq. (F1)

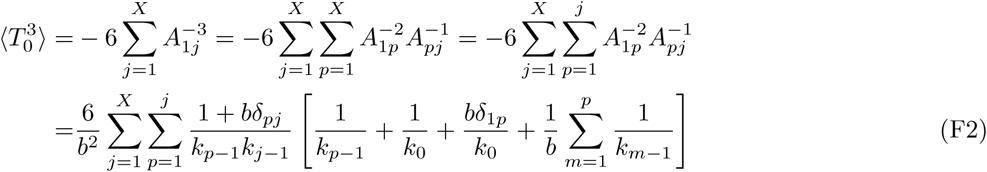

Next we define a vector ⟨**T** ⟩^4^, whose components are fourth moments of the FPTs starting from various protein counts *i*. Again doing integration by parts successively, we get:

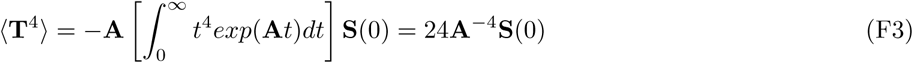

So,

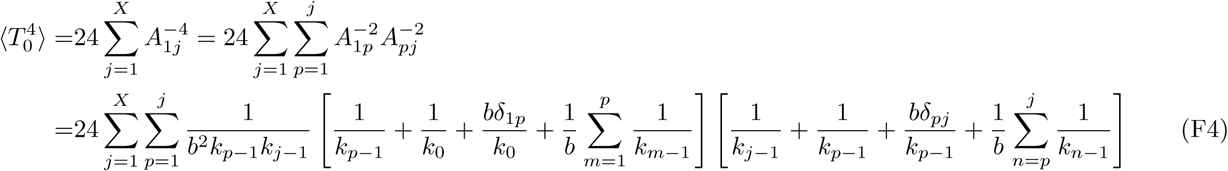

## Appendix G Alternative formalism to calculate FPT distribution for general case of unequal transition rates with no degradation

Here we begin with Eq. (7) in the main text and evaluate *exp*(**A***t*).

Firstly we note from Eq. (5) in main text that matrix **A** is upper triangular, for *γ* = 0 and hence the eigenvalues are given by its diagonal elements:

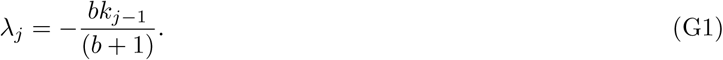

Next we show that the matrix which diagonalises **A** such that **D**^*−***1**^**AD** = **A**_**D**_ (with (**A**_**D**_)_*ij*_ = *λ*_*j*_*δ*_*ij*_) has its elements given by:

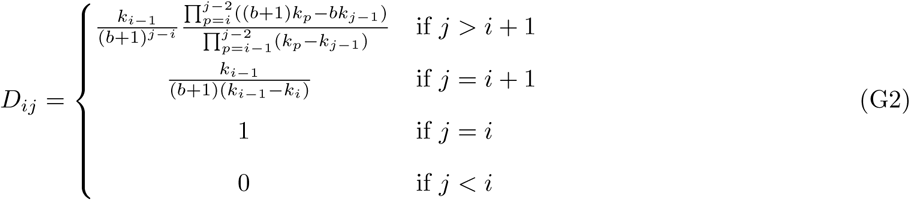

With the above form of **D** it may be shown directly that

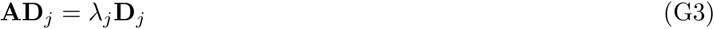

such that the vector **D**_*j*_ is the *j*-th column of **D** and an eigenvector of **A**.

**Proof of AD**_*j*_ = *λ*_*j*_**D**_*j*_:

Using Eqs (5) (in main text), (G1) and (G2), we get for *j >* (*i* + 1),

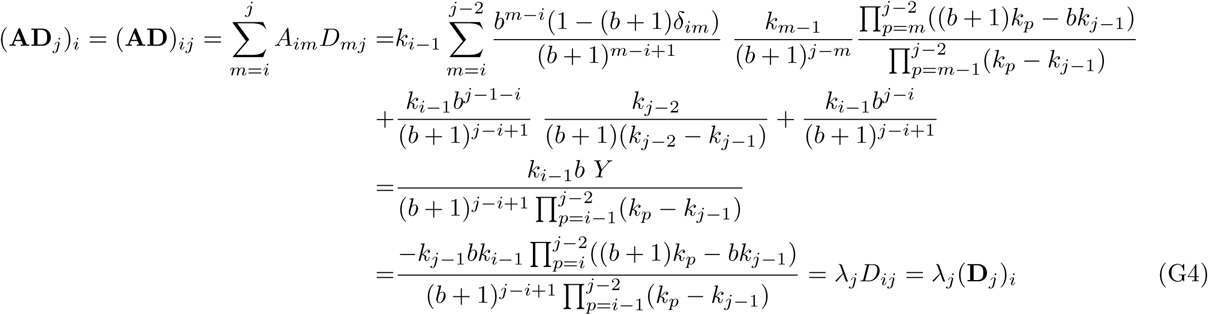

The quantity *Y* in the intermediate state above is a sum of several terms as follows, which simplify on repeated factorisation and cancellation to give the short expression in the last step above:

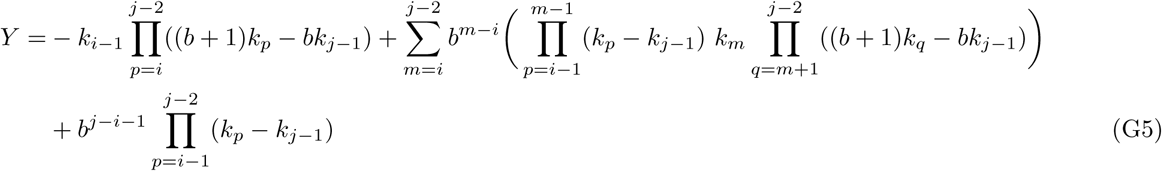

In the special case of j = i + 1,

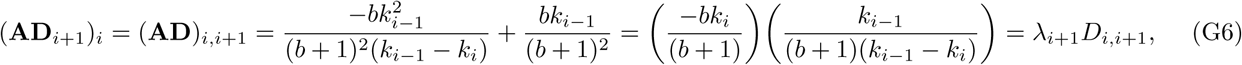

and for *j* = *i*

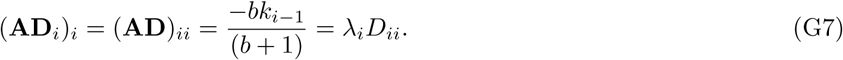

Hence, proved.

The elements of the inverse matrix **D**^*−***1**^ is given by

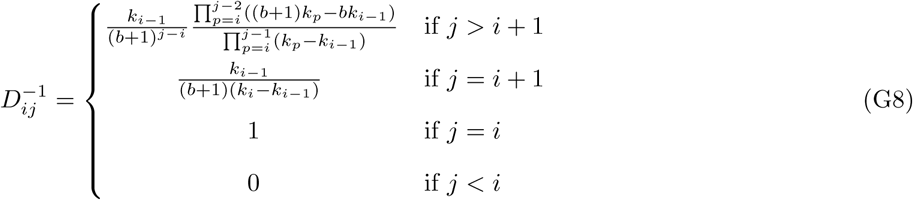

Using the above diagonal form, the vector of different first passage probability distributions in Eq. (7) may be written as

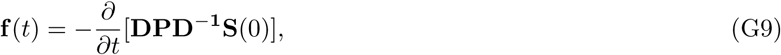

where **P** = exp(**A**_**D**_*t*) = **D**^*−***1**^ exp(**A***t*)**D**. Thus (**P**)_*mn*_ = exp(*λ*_*m*_*t*)*δ*_*mn*_. Moreover we are interested in the case of initial protein number being 0, and in the corresponding first passage distribution

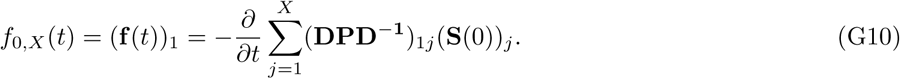

From Eq. (G8), 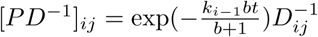, and then it follows from Eq.(G2) that

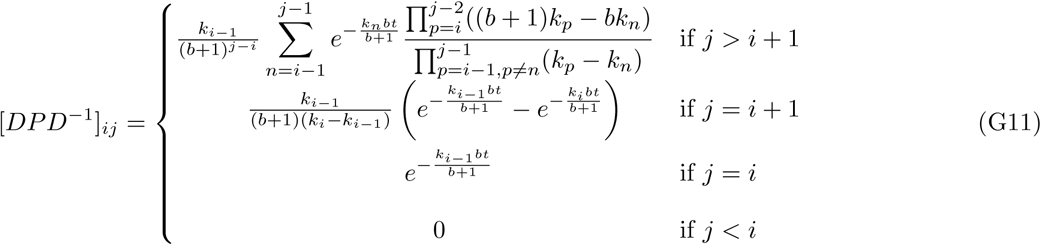

Recalling that all the elements of **S**(0) are 1, we have from Eq. (G10),

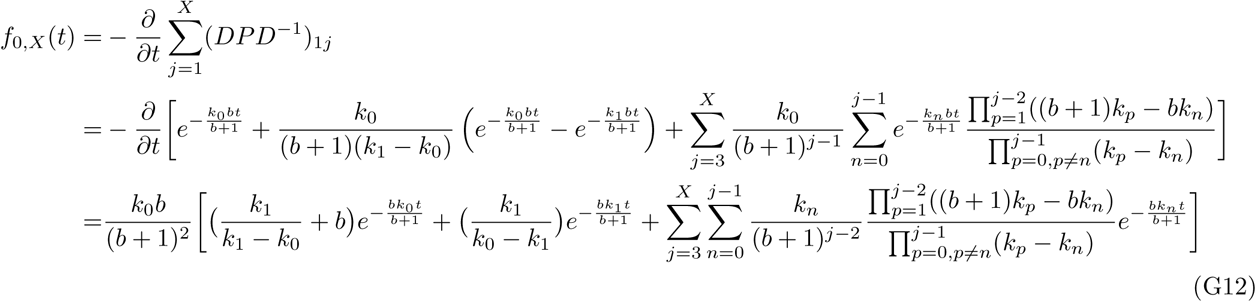

The above expression is valid for *X >* 2, and for the special cases of *X* = 1,

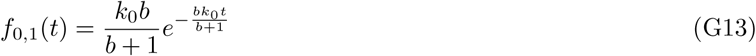

and *X* = 2,

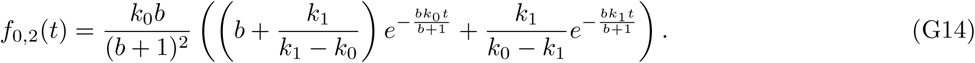

## Appendix H Alternative formalism to calculate FPT distribution for special case of equal transition rates with no degradation

As mentioned in the main text, we can take proper limit of Eq. (10) to get non divergent FPT expression for this special case. We first write the first passage distributions for *X* = 1, 2, 3, 4 below in a particular way to observe a pattern:

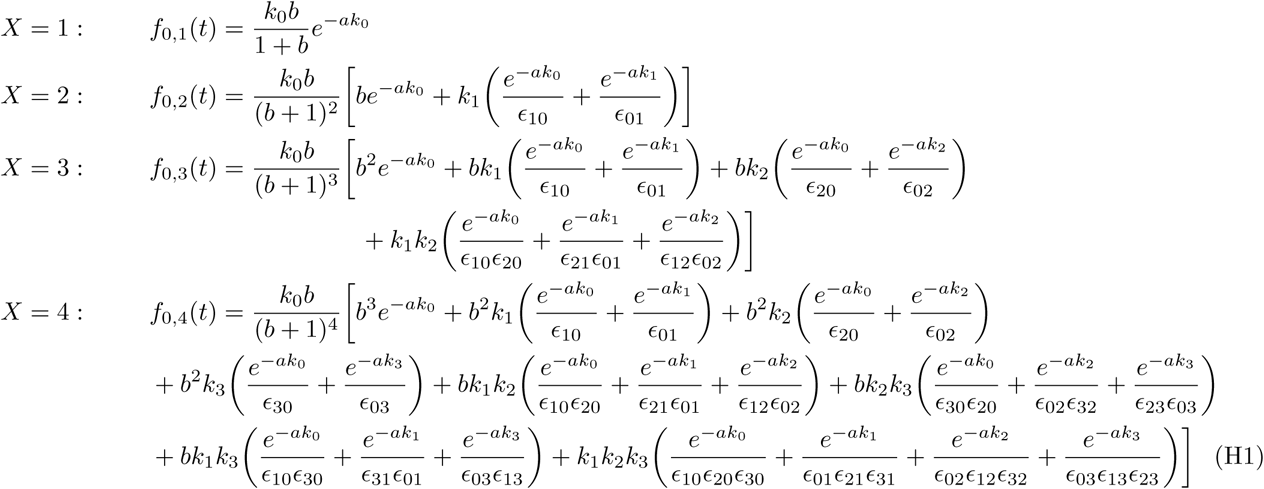

In the above *a* = *bt/*(*b* + 1), and *E*_*ij*_ = (*k*_*i*_ *− k*_*j*_). Note that in Eqs. (H1), every coefficient within parentheses (*…*) may be represented by a symbo 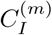 involving a certain set of integers *I* = {0, *i*_1_, *i*_2_,, *i*_*m*_}. The general form of the coefficient is as follows:

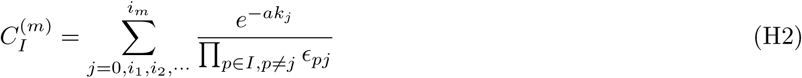

The coefficients 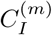 have prefactors 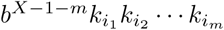, and terms corresponding to all possible 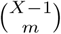 combinations (*comb*) of *m* integers out of the set *{*1, 2, *…, X −* 1*}* are added. Thus the general form of *f*_0,*X*_ (*t*), with 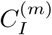 from Eq. (H2), is finally:

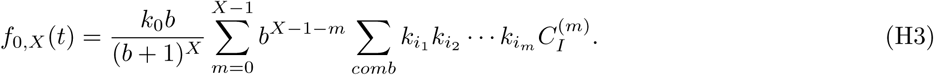

We then take the limit that *k*_*j*_ for all *j* approach the same value *k* by first writing *k*_*j*_ = *k* +Δ*j* and taking *lim*Δ *→* 0. By writing the exponential in the numerator of Eq. (H2) in series form, and noting all terms for *l > m* vanish with Δ *→* 0, we have 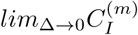 given by:

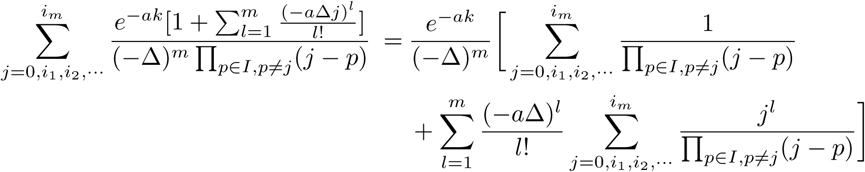

At this point we assume the following two identities (see the discussion on proofs of these in Appendix I below):

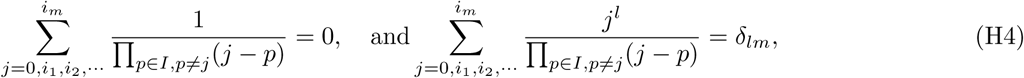

Using Eqs. (H4) we have

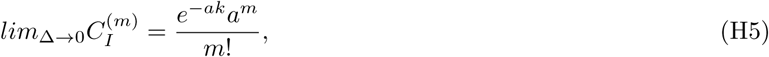

and as 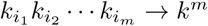 in the same limit, we have from Eq. (H3),

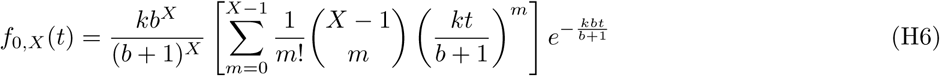

The term inside the square bracket is a confluent hypergeometric function _1_*F*_1_, and hence

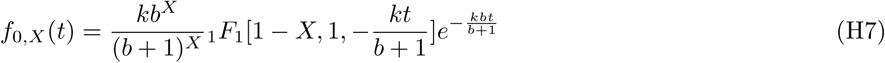

## Appendix I Discussion on proof of the identities in Eqs. (H4)

A proof can be give for a special case first. Let the set *I* be a simple set, *I* = {0, 1, 2, *…, m*} and 1 *≤ j ≤ m* It is easy to see that

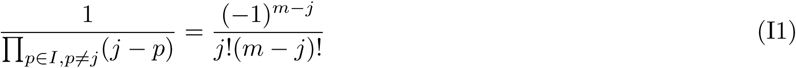

Using Eq. (I1),

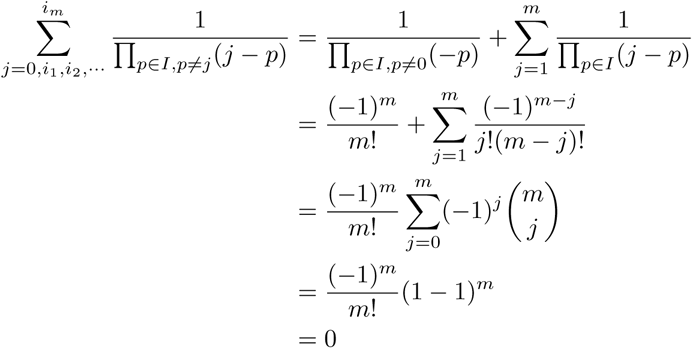

Hence, the first part of Eq. (H4) is proved, for the simple case. The second part is

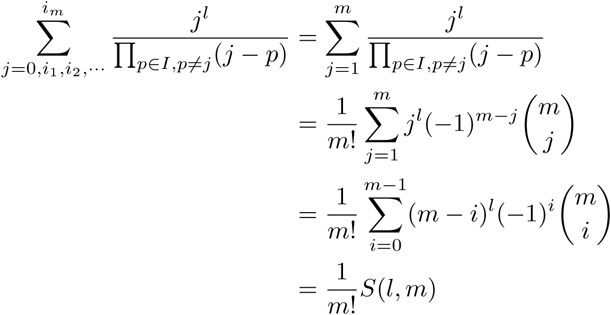

*S*(*l, m*) is known to be the number of surjective maps from a set with *l* elements to another set with *m* elements. Only when *l* is equal to *m*, surjective mapping is possible. The number of surjective maps is then equal to the number of ways of putting *m* distinct balls into *m* distinct boxes (no box being empty) =*m*!*δ*_*lm*_. Hence, 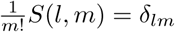 and the identity for the special case is proved.

For the general case, we conjecture that the identity is valid for any random set of *m* positive integers. Although we cannot give a proof, we computationally checked it using Mathematica. The set of integers *I* = {0, *i*_1_, *i*_2_,, *i*_*m*_} were chosen randomly 1000 times. For every such set, the identity was verified. We varied *m* such that 1 *≤ m ≤* 100. In that way, the identity was checked 1000 *×* 100 = 10^5^ times, for the general case. A formal proof would be nice to find in future.

